# Choice of library preparation and its effects on sequence quality, genome assembly, and precise *in silico* prediction of virulence genes in shiga toxin producing *Escherichia coli*

**DOI:** 10.1101/2020.11.02.364646

**Authors:** Julie Haendiges, Karen Jinneman, Narjol Gonzalez-Escalona

**Affiliations:** Division of Microbiology, Office of Regulatory Science; Center for Food Safety and Applied Nutrition, Food and Drug Administration, College Park, MD; Pacific Regional Laboratory NW, Office of Regulatory Affairs, US Food and Drug Administration, Bothell, WA

**Author notes:** Corresponding authors. and. Mailing address, Center for Food Safety and Applied Nutrition, Food and Drug Administration, 5001 Campus Drive, College Park, MD 20740, USA.

**Keywords:** *Escherichia coli*, MiSeq sequencing, foodborne pathogen, flour, DNA Prep, Nextera XT

## Abstract

Whole genome sequencing (WGS) provides essential public health information and is used worldwide for pathogen surveillance, epidemiology, and source tracking. The sequencing of foodborne pathogens is commonly performed with Illumina sequencing chemistry to obtain data with high accuracy. The choice of library preparation method for highly complex organisms is very critical and can affect the final data output. The majority of Illumina sequencing platforms use rapid library preparation such as Nextera XT (transposon-based technology) (Illumina San Diego, CA), but this preparation has the potential to miss randomly distributed segments of genomes that might be important for downstream analyses. The Illumina Nextera DNA Prep library preparation kit, the successor of Nextera XT, shows better overall coverage of the complete genome. This study compared the quality of sequence data generated using Nextera XT and Nextera DNA Prep kits for DNA library preparation on an Illumina MiSeq, using a set of 30 O121:H19 shiga-toxin positive *Escherichia coli* strains isolated from flour during a 2016 outbreak. The performance of the two kits were evaluated using several metrics including sequencing quality, assembly quality, uniformity of genome coverage, and virulence gene identification. Overall, the results showed that in all of the analysed metrics, the Nextera DNA Prep kit performed outstanding in comparison to Nextera XT. The Nextera DNA Prep kit allowed for comprehensive detection of all virulence genes, which is of extremely high importance for making an educated assessment of the virulence potential of *Escherichia coli*. This comprehensive side-by-side comparison will be of significance for those interested in improving their sequencing workflow for STECs and the determination of health risks using WGS data.

## Introduction

Whole genome sequencing (WGS) has been used in public health surveillance and outbreak detection of foodborne illnesses since 2012 (1). WGS not only allows for phylogenetic analysis of strains but also for determination of serotype, virulence factors, and antimicrobial resistance. The majority of the bacterial whole genome sequencing effort has been conducted utilizing Illumina Nextera XT (Illumina, San Diego, CA) reagents for library preparation, which utilizes enzymatic fragmentation using transposon enzymes (2). Overall, Nextera XT is effective for producing sequences that are optimal for determining phylogenies and source tracking. However, this method has been shown previously to have representative bias in certain regions of the genomes, creating the potential to overlook randomly distributed segments that could be important for downstream analyses (3-5). Organisms such as shiga-toxin positive *Escherichia coli* (STEC), present a unique problem as they possess very complex genomes with repetitive elements (i.e. insertion sequences, phages and plasmids) (5-7), and the choice of DNA library preparation can affect the final output. Therefore, there is the need for a method that results in better coverage of the entire genome.

The classification of pathogenic *E. coli* (EPEC, ETEC, EHEC, EAEC) depends on the presence of pathogenicity islands, virulence genes, and plasmids that cause diarrheal illness. In order for an *E. coli* strain to be considered a STEC and cause human illness, the strain must contain genes that allow attachment, colonization, and production of Shiga toxin (8-11). The majority of virulence genes are found in the chromosome, but there are also those contained on the virulence plasmid (8, 10, 12). All STECs contain a virulence plasmid, that may differ in gene content, and this assortment of virulence genes can affect clinical outcomes and are of high importance for surveillance (13, 14). This highlights the importance of producing a library that contains the complete STEC genome in order to make an informed decision about the public health impact of the analyzed strain.

A recent DNA library kit released by Illumina, Nextera DNA Prep, shows overall better coverage of the complete genome. There are a few differences between these kits, including the fragmentation process of the transposome, dual size selection and the quality of the indices (2). The Nextera DNA Prep kit, previously known as the Nextera DNA Flex Library Prep, utilizes a magnetic-bead linked transposome (BLT) with a known fixed concentration of enzyme (2). According to Bruinsma et al. (2019), the efficiency of fragmentation is improved due to the consistent enzyme to DNA ratio (2). The fixed quantity of transposome on the bead is saturated at high DNA input concentrations (>100ng) and allows for more uniform insert size and concentration of the final libraries. The DNA Prep kit has previously been shown to work very well with organisms of variable GC content and results in an even distribution of read depth across the genome unlike XT (4). Library preparation time is similar to XT, even with addition of a dual size selection step. The libraries are self-normalized when using high DNA inputs (>100 ng) and the concentration of input DNA is less critical, which saves time during the initial quantification and dilutions.

This study evaluated the genomic sequence data generated from library preparations using the XT and DNA Prep kits. The target genes examined in the study included Antimicrobial Resistance (AMR) genes, virulence genes, and plasmid identification for STEC characterization. Library preparation is a critical and rate-limiting step (4) therefore a method that achieves better genome coverage coupled with a reliable and fast protocol is necessary to accomplish accurate determination of virulence potential. The performance of both transposon-based library preparation kits (XT and DNA Prep) was determined through a characterization comparison of the libraries from a subset of STEC strains isolated during a 2016 multistate outbreak associated with contaminated flour (15). The library preparation methods were evaluated based on the following metrics: assembly quality, *in silico* determination of serotype, MLST, AMR genes, and virulence genes, as well as single nucleotide polymorphism (SNP) phylogenetic analysis.

## Materials and methods

### Bacterial strains and DNA preparation

The strains were isolated from various types of flour by the U.S. Food and Drug Administration ORA Pacific Regional laboratory as part of a Federal public health multistate *Escherichia coli* investigation. The isolates were grown overnight in trypticase soy broth (TSB) (Becton, Dickinson and Company, Franklin Lakes, NJ) at 37°C and genomic DNA was extracted using the Qiagen DNeasy Blood and Tissue Kit (Qiagen, Valencia, CA). The concentration of resultant DNA was determined using a Qubit double-stranded DNA BR assay kit and a Qubit fluorometer (ThermoFisher Scientific, Waltham, MA), according to the manufacturer’s instructions, then stored at -20°C until use. The 30 shiga-toxin positive strains used in this study are listed in Table 1.

**Table 1.** Metadata for the 30 STEC strains used in this study.

| CFSAN No. | Isolate name | Source | Location | Date of Collection | Serotype | Biosample Accession | ST<sup>a</sup> |
| --- | --- | --- | --- | --- | --- | --- | --- |
| CFSAN051559 | FNW19M89 | All-Purpose Flour | USA | 06/2016 | O121:H19 | SAMN05245391 | 655 |
| CFSAN051560 | FNW19M90 | All-Purpose Flour | USA | 06/2016 | O121:H19 | SAMN05245392 | 655 |
| CFSAN051561 | FNW19M91 | All-Purpose Flour | USA | 06/2016 | O121:H19 | SAMN05245393 | 655 |
| CFSAN051562 | FNW19M92 | All-Purpose Flour | USA | 06/2016 | O121:H19 | SAMN05245394 | 655 |
| CFSAN051563 | FNW19M93 | All-Purpose Flour | USA | 06/2016 | O121:H19 | SAMN05245395 | 655 |
| CFSAN051564 | FNW19M94 | All-Purpose Flour | USA | 06/2016 | O121:H19 | SAMN05245396 | 655 |
| CFSAN051565 | FNW19M95 | All-Purpose Flour | USA | 06/2016 | O121:H19 | SAMN05245618 | 655 |
| CFSAN051566 | FNW19M96 | All-Purpose Flour | USA | 06/2016 | O121:H19 | SAMN05245621 | 655 |
| CFSAN051567 | FNW19M97 | All-Purpose Flour | USA | 06/2016 | O121:H19 | SAMN05245624 | 655 |
| CFSAN051568 | FNW19M98 | All-Purpose Flour | USA | 06/2016 | O121:H19 | SAMN05245744 | 655 |
| CFSAN051758 | FNW19N01 | Unbleached White Flour | USA:MI | 06/2016 | O121:H19 | SAMN05289759 | 655 |
| CFSAN051759 | FNW19N02 | Unbleached White Flour | USA:MI | 06/2016 | O121:H19 | SAMN05289761 | 655 |
| CFSAN051760 | FNW19N03 | Unbleached White Flour | USA:MI | 06/2016 | O121:H19 | SAMN05289763 | 655 |
| CFSAN051761 | FNW19N04 | Unbleached White Flour | USA:MI | 06/2016 | O121:H19 | SAMN05289764 | 655 |
| CFSAN051762 | FNW19N05 | Unbleached White Flour | USA:MI | 06/2016 | O121:H19 | SAMN05289767 | 655 |
| CFSAN051763 | FNW19N06 | Unbleached White Flour | USA:MI | 06/2016 | O121:H19 | SAMN05289769 | 655 |
| CFSAN051764 | FNW19N07 | Unbleached White Flour | USA:MI | 06/2016 | O121:H19 | SAMN05289771 | 655 |
| CFSAN051765 | FNW19N08 | Unbleached White Flour | USA:MI | 06/2016 | O121:H19 | SAMN05289773 | 655 |
| CFSAN051766 | FNW19N09 | Unbleached White Flour | USA:MI | 06/2016 | O121:H19 | SAMN05289774 | 655 |
| CFSAN051767 | FNW19N10 | Unbleached White Flour | USA:MI | 06/2016 | O121:H19 | SAMN05289776 | 655 |
| CFSAN051768 | FNW19N11 | Unbleached White Flour | USA:MI | 06/2016 | O121:H19 | SAMN05289777 | 655 |
| CFSAN051769 | FNW19N12 | Unbleached White Flour | USA:MI | 06/2016 | O121:H19 | SAMN05289778 | 655 |
| CFSAN051770 | FNW19N13 | Unbleached White Flour | USA:MI | 06/2016 | O121:H19 | SAMN05289779 | 655 |
| CFSAN051771 | FNW19N14 | Unbleached White Flour | USA:MI | 06/2016 | O121:H19 | SAMN05289780 | 655 |
| CFSAN051772 | FNW19N15 | Unbleached White Flour | USA:MI | 06/2016 | O121:H19 | SAMN05289781 | 655 |
| CFSAN052204 | FNW19N17 | Enriched White Flour | USA:OK | 06/2016 | O121:H19 | SAMN05294031 | 655 |
| CFSAN052205 | FNW19N18 | Enriched White Flour | USA:OK | 06/2016 | O121:H19 | SAMN05294032 | 655 |
| CFSAN052206 | FNW19N19 | Enriched White Flour | USA:OK | 06/2016 | O121:H19 | SAMN05294033 | 655 |
| CFSAN052207 | FNW19N20 | Enriched White Flour | USA:OK | 06/2016 | O121:H19 | SAMN05294034 | 655 |
| CFSAN052208 | FNW19N21 | Enriched White Flour | USA:OK | 06/2016 | O121:H19 | SAMN05294035 | 655 |

<sup>a</sup> http://enterobase.warwick.ac.uk/species/index/ecoli.

### Closure of reference genome using MinION and Flye

For further bioinformatic analysis a closed reference strain was necessary. Strain FNW19M81 was obtained and grown overnight in TSB at 37°C. The genomic DNA was isolated using the Maxwell RSC Cultured Cells DNA kit (Promega, Madison, WI) following the manufacturer’s protocols with the addition of RNase A (ThermoFisher Scientific, Waltham, MA) treatment. Closure of this genome was performed using both short and long read sequencing technology. The short-read sequencing library was prepared using Illumina DNA Prep and the long-read sequencing library was prepared using the Ligation Sequencing Kit (SQK-LSK108) (Oxford Nanopore Technologies, Oxford, UK). Short read sequencing was performed using Illumina v3 sequencing reagents on an Illumina MiSeq while the long reads were sequenced on a FLO-MIN106 (R9.4) flowcell for 48 hours on an Oxford Nanopore MinION device. The long-reads were live basecalled using Albacore v2.0.1 included in the MinKNOW software (v1.10.11). All reads below 5,000 basepairs in length were removed from further analysis. The assembly procedure was performed as described in (16) with the following alterations: the assembly of the long reads was performed with Flye v1.6 (17) instead of Canu (18) and for the hybrid assembly Unicycler v0.4.8 (19) was used instead of SPAdes (20). The genome was deposited in GenBank under accession number CP051631 and CP051632. The final assembly of the chromosome and plasmid were annotated using Prokka v1.13 (21).

### Library preparation and whole genome sequencing

In order to compare the data generated by the two library preparation kits, the same DNA extract was used as input for both library preparations. The DNA library preparations were performed according to published protocols by Illumina (Illumina, Nextera DNA Prep Library Prep Reference Guide) and FDA GenomeTrakr network (Illumina, Nextera XT Library Prep Reference Guide). The XT kit uses 1 ng of DNA input and the DNA Prep kit allows for DNA input from 1-500 ng. For this study we used the highest input of DNA possible with the remaining volume for library preparation with the DNA Flex kit. The DNA input value for each strain can be found in Table 2. An Illumina MiSeq benchtop sequencer (Illumina, San Diego, CA) was used to sequence the two library preparations with v3 sequencing chemistry with 2×250 bp pair-end reads.

**Table 2.** Assembly statistics by DNA library kit for each strain. (PREP – Nextera DNA Prep, XT – Nextera XT)

| Strain | DNA library prep | SRA Accession | Total DNA Input (ng) | Genome coverage X | > Q30 Avg. Read lengths | Average insert size | # Contigs >500 bp | Largest Contig (bp) | N50 (bp) |
| --- | --- | --- | --- | --- | --- | --- | --- | --- | --- |
| CFSAN051559 | XT | SRR3747656 | 1 | 108 | 160 | 260 | 233 | 468,170 | 124,862 |
|  | PREP | SRR11507285 | 13 | 98 | 226 | 304 | 230 | 468,170 | 134,860 |
| CFSAN051560 | XT | SRR3747657 | 1 | 70 | 162 | 265 | 279 | 308,556 | 90,323 |
|  | PREP | SRR11508023 | 12 | 112 | 231 | 323 | 225 | 468,170 | 149,318 |
| CFSAN051561 | XT | SRR3747658 | 1 | 37 | 166 | 243 | 250 | 308,413 | 122,675 |
|  | PREP | SRR11507295 | 12 | 111 | 231 | 328 | 223 | 468,170 | 149,318 |
| CFSAN051562 | XT | SRR3747659 | 1 | 32 | 166 | 262 | 250 | 468,170 | 122,457 |
|  | PREP | SRR11507316 | 83 | 92 | 238 | 359 | 213 | 468,170 | 161,492 |
| CFSAN051563 | XT | SRR3747660 | 1 | 121 | 164 | 251 | 267 | 468,170 | 114,952 |
|  | PREP | SRR11507718 | 17 | 97 | 233 | 326 | 226 | 468,251 | 136,153 |
| CFSAN051564 | XT | SRR3747661 | 1 | 59 | 169 | 265 | 231 | 468,245 | 122,671 |
|  | PREP | SRR11507313 | 93 | 118 | 236 | 327 | 203 | 468,170 | 161,492 |
| CFSAN051565 | XT | SRR3747662 | 1 | 32 | 159 | 266 | 229 | 468,855 | 118,199 |
|  | PREP | SRR11507300 | 81 | 101 | 237 | 339 | 247 | 468,170 | 136,153 |
| CFSAN051566 | XT | SRR3747677 | 1 | 81 | 166 | 271 | 232 | 468,169 | 134,860 |
|  | PREP | SRR11507286 | 14 | 124 | 232 | 333 | 225 | 468,251 | 136,153 |
| CFSAN051567 | XT | SRR3747663 | 1 | 173 | 168 | 245 | 222 | 468,855 | 134,860 |
|  | PREP | SRR11507293 | 13 | 102 | 233 | 324 | 212 | 468,170 | 136,153 |
| CFSAN051568 | XT | SRR3747664 | 1 | 74 | 158 | 253 | 232 | 371,303 | 92,524 |
|  | PREP | SRR11506703 | 124 | 101 | 239 | 343 | 192 | 468,170 | 161,492 |
| CFSAN051758 | XT | SRR3713425 | 1 | 107 | 189 | 251 | 245 | 468,251 | 134,860 |
|  | PREP | SRR11507309 | 11 | 131 | 228 | 320 | 230 | 468,251 | 136,153 |
| CFSAN051759 | XT | SRR3713426 | 1 | 109 | 195 | 268 | 218 | 468,855 | 124,722 |
|  | PREP | SRR11507292 | 12 | 99 | 230 | 326 | 214 | 468,170 | 136,153 |
| CFSAN051760 | XT | SRR3713427 | 1 | 123 | 166 | 257 | 218 | 468,170 | 136,153 |
|  | PREP | SRR11507315 | 11 | 98 | 227 | 323 | 212 | 468,170 | 149,318 |
| CFSAN051761 | XT | SRR3713429 | 1 | 113 | 192 | 236 | 272 | 462,510 | 118,379 |
|  | PREP | SRR11507717 | 122 | 104 | 239 | 347 | 211 | 468,170 | 134,860 |
| CFSAN051762 | XT | SRR3713430 | 1 | 115 | 168 | 250 | 261 | 398,462 | 111,929 |
|  | PREP | SRR11647983 | 27 | 121 | 226 | 322 | 205 | 468,170 | 136,153 |
| CFSAN051763 | XT | SRR3713431 | 1 | 72 | 230 | 331 | 260 | 277,234 | 100,914 |
|  | PREP | SRR11508014 | 101 | 109 | 239 | 343 | 214 | 468,170 | 136,153 |
| CFSAN051764 | XT | SRR3717432 | 1 | 117 | 220 | 295 | 212 | 468,170 | 147,488 |
|  | PREP | SRR11647979 | 12 | 86 | 223 | 310 | 210 | 468,170 | 136,153 |
| CFSAN051765 | XT | SRR3713433 | 1 | 90 | 229 | 354 | 225 | 468,170 | 147,488 |
|  | PREP | SRR11508027 | 81 | 98 | 237 | 341 | 236 | 468,170 | 161,492 |
| CFSAN051766 | XT | SRR3713434 | 1 | 112 | 183 | 270 | 224 | 468,170 | 122,671 |
|  | PREP | SRR11507380 | 112 | 113 | 239 | 343 | 210 | 468,170 | 136,153 |
| CFSAN051767 | XT | SRR3713435 | 1 | 107 | 169 | 242 | 222 | 468,170 | 134,860 |
|  | PREP | SRR11508026 | 105 | 121 | 240 | 362 | 203 | 468,170 | 149,318 |
| CFSAN051768 | XT | SRR3713436 | 1 | 90 | 205 | 281 | 221 | 468,170 | 134,860 |
|  | PREP | SRR11507722 | 78 | 98 | 239 | 348 | 213 | 468,170 | 136,153 |
| CFSAN051769 | XT | SRR3713563 | 1 | 108 | 197 | 266 | 244 | 468,251 | 132,264 |
|  | PREP | SRR11507308 | 82 | 94 | 240 | 357 | 219 | 468,170 | 136,153 |
| CFSAN051770 | XT | SRR3713437 | 1 | 110 | 182 | 261 | 224 | 468,170 | 134,860 |
|  | PREP | SRR11506716 | 69 | 90 | 239 | 360 | 215 | 468,170 | 149,318 |
| CFSAN051771 | XT | SRR3713438 | 1 | 111 | 221 | 312 | 241 | 468,170 | 147,488 |
|  | PREP | SRR11507290 | 30 | 103 | 236 | 328 | 243 | 468,170 | 136,153 |
| CFSAN051772 | XT | SRR3713439 | 1 | 105 | 198 | 269 | 232 | 468,936 | 134,860 |
|  | PREP | SRR11507381 | 66 | 94 | 239 | 348 | 223 | 468,170 | 161,492 |
| CFSAN052204 | XT | SRR3743154 | 1 | 101 | 180 | 259 | 238 | 468,170 | 134,860 |
|  | PREP | SRR11507299 | 70 | 97 | 239 | 361 | 214 | 468,170 | 136,153 |
| CFSAN052205 | XT | SRR3743156 | 1 | 89 | 177 | 266 | 231 | 468,855 | 134,860 |
|  | PREP | SRR11507282 | 59 | 94 | 238 | 361 | 217 | 468,170 | 136,153 |
| CFSAN052206 | XT | SRR3743157 | 1 | 165 | 192 | 271 | 231 | 468,251 | 134,860 |
|  | PREP | SRR11507724 | 72 | 86 | 240 | 370 | 213 | 468,170 | 161,492 |
| CFSAN052207 | XT | SRR3743158 | 1 | 116 | 203 | 266 | 243 | 468,855 | 134,860 |
|  | PREP | SRR11647986 | 88 | 96 | 240 | 357 | 207 | 468,170 | 161,492 |
| CFSAN052208 | XT | SRR3743159 | 1 | 111 | 193 | 266 | 225 | 468,855 | 134,860 |
|  | PREP | SRR11647987 | 149 | 111 | 240 | 352 | 209 | 348,419 | 136,153 |

### Bioinformatic analysis

Two *de novo* assemblies were generated from the reads of each isolate using SPAdes v3.13.0 (20), using k-mer lengths of [21, 33, 55, 77, 99, 127], and the options --careful and --only-assembler. All assemblies were quality checked using Quast v5.0 (22) and evaluated for the number of contigs, largest contig length, and N50. All contigs with a length below 500 bp were trimmed from the final assemblies before proceeding to analysis with AMRfinder.

The initial analysis and identification of the strains were performed using an *in silico E. coli* MLST approach, based on the information available at the *E. coli* MLST website (http://mlst.warwick.ac.uk/mlst/dbs/Ecoli) and using Ridom SeqSphere+ software v2.4.0 (Ridom; Münster, Germany) (http://www.ridom.com/seqsphere). Seven housekeeping genes (*adk, fumC, gyrB, icd, mdh, recA*, and *purA*), described previously for *E. coli* (23), were used for MLST analysis. The same *E. coli* MLST database was also used to assign numbers for alleles and sequence types (STs). The serotype of each strain analyzed in this study was confirmed using the genes deposited in the Center for Genomic Epidemiology (http://www.genomicepidemiology.org) for *E. coli* as part of their web-based serotyping tool (SerotypeFinder 1.1 https://cge.cbs.dtu.dk/services/SerotypeFinder) (24). Each whole genome sequence was screened for O-type or H-type genes.

Virulence genes, stress genes, and antimicrobial resistance (AMR) genes were identified using the AMRFinder v3.6.10 command line tool (25). All assemblies were analyzed using the --plus option to include *E. coli* virulence genes and stress tolerance genes. This database has over 600 virulence reference sequences, 200 stress reference sequences, and 6,000 AMR reference sequences. The virulence genes included in the database represent a repertoire of genes found in different *E. coli* pathotypes (ETEC, STEC, EAEC, and EPEC) in order to detect any possible *E. coli* hybrid present (26).

### Phylogenetic analysis

The phylogenetic relationship of the strains was assessed by the CFSAN SNP Pipeline (27). The closed genome of FNW19M81 was used as the reference for analysis. In addition to the 30 strains in this study, three clinical strains not associated with the outbreak were used for analysis to serve as outgroups. To evaluate if Single Nucleotide Polymorphism (SNP) calls were affected by library preparation, the raw reads from both library preparations, for all strains, was analyzed. A maximum likelihood tree was generated using RAxML (28) from the SNP matrix, using 500 bootstraps and the GTRCAT substitution model to identify the best tree.

### Nucleotide sequence accession numbers

The draft genome sequences for all 30 STEC strains used in our analyses are available in GenBank under the accession numbers listed in Table 1.

## Results

### Reference strain analysis

In order to accurately assess the differences in performance of both DNA library preparations we closed one representative of the outbreak strains (strain FNW19M81). The complete closed circular genome for this O121:H19 strain resulted in a genome with one chromosome (CP051631) of length 5,391,339 bp (50.7% GC) and a single plasmid (CP051632) of length 81,965 bp (46.3% GC). This strain was used to test for inclusivity of all virulence, AMR, and stress tolerance genes. Analysis of the genome of this strain identified the presence of 19 virulence genes with three of them: *ehxA, toxB*, and *espP* located on the plasmid.

### Evaluation of assembly quality

To directly compare the results and quality of the different library preparations the same DNA extract was used by both library methods. Two *de novo* assemblies were produced for each of the thirty strains, one for each library preparation (XT and DNA Prep). The quality of these assemblies was evaluated by examining the Quast and SPAdes outputs, focusing mainly on the values of contigs above >500 bp, largest contig, N50, average insert size, and read length quality. These results are shown in Table 2. The majority of the assemblies from the DNA Prep library preparations show fewer number of contigs (n=27/30) and larger N50 (n= 28/30) as seen in Figure 1A and B. The assemblies from XT ranged from 203 to 279 contigs while the assemblies from DNA Prep ranged from 192 to 247 total contigs.

**Figure 1.**
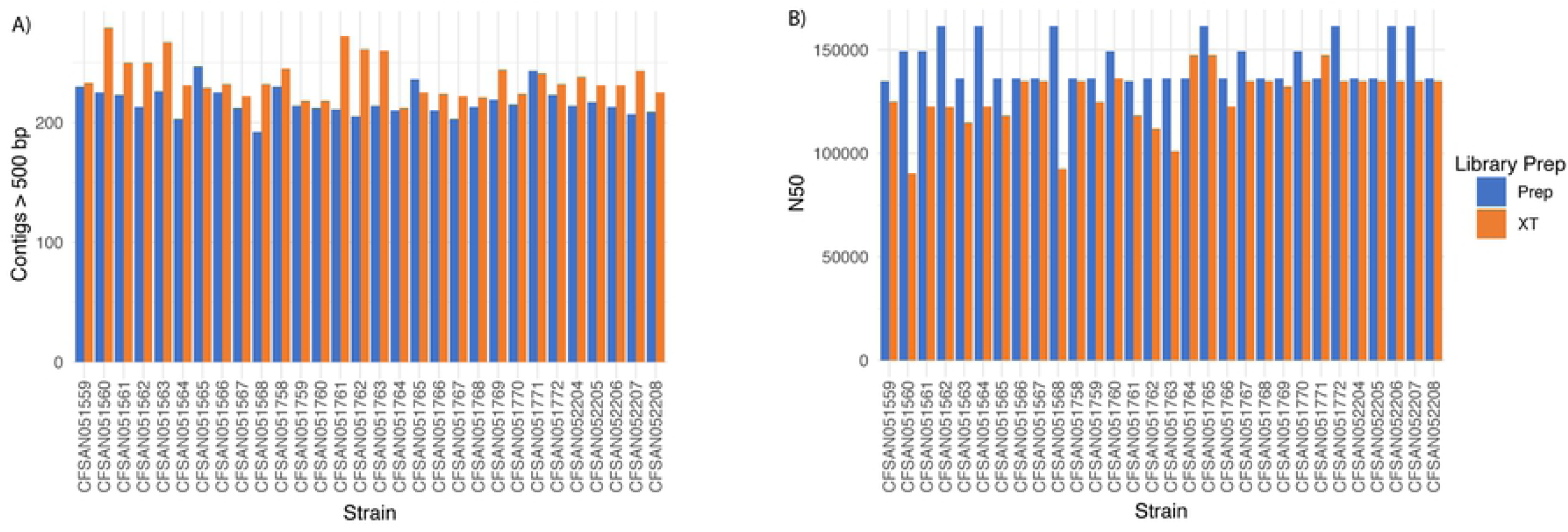
Comparison of the contigs and N50 obtained per DNA library kit and strain. The DNA Prep kit resulted in sequencing reactions that rendered the best assemblies and N50s per strains. A) Number of contigs larger than 500 basepairs per DNA library kit and strain. B) N50 for the assemblies by DNA library kit and strain.

The average read lengths at or above Q30 from DNA Prep were 234 bp out of 250 bp, while the reads from XT were 184 bp out of 250 bp. The average DNA insert size with DNA Prep was 335 bp (ranging from 304 to 370bp) compared to libraries from XT with an average insert size of 260 bp (range 242-354) allowing for less overlap of read 1 and read 2 (4). Figure 2 shows the distribution of paired read length (insert sizes) for the two library preparations for strain CFSAN051560. The distribution of paired read length (insert sizes) of DNA Prep libraries was consistent across all of the libraries similar to CFSAN051560 while those from XT show a higher concentration of smaller insert sizes (data not shown).

**Figure 2.**
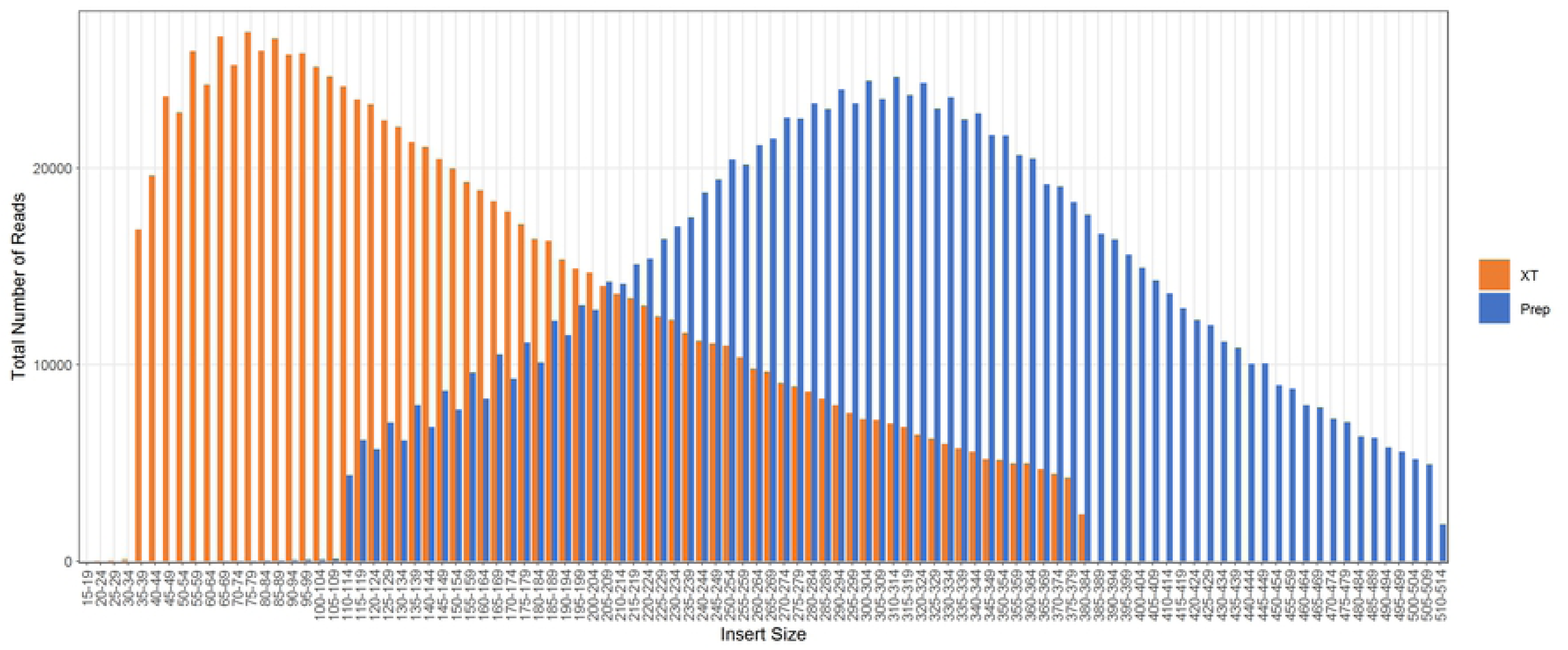
The distribution of the total number of reads based on the size of the paired read length (insert size) for strain CFSAN051560. DNA Prep results are shown in blue and Nextera XT is shown in orange.

### *In silico* MLST, molecular serotyping and AMRFinder results

*In silico* MLST and molecular serotyping showed that all strains belonged to ST655 and O121:H19, respectively, using both DNA library preparation kits. Assemblies from both library preparations for each sample were interrogated using AMRFinder plus (25) to identify the presence of *E. coli* virulence, AMR, and stress tolerance genes. The results are shown in Table 3. All strains from this outbreak contained the following AMR genes: *acrF, blaEC*, and *mdtM* and the following stress genes: *terD, terW, terZ*, and *ymgB*. These genes were all identified equally by both library preparations.

**Table 3.** AMRFinder Results for the 30 STEC strains used in this study by DNA library preparation kit.

| Strains | DNA<br>library<br>prep | *efal* | *espF* | *espI* | *nleC* | *tccP* | *ehxA* | *espP* | *toxB* | PCR-Plasmid<sup>a</sup> |
| --- | --- | --- | --- | --- | --- | --- | --- | --- | --- | --- |
| FNW19M81<br>Reference |  | Y | Y | Y | Y | Y | Y | Y | Y | + |
| CFSAN051560 | XT | Y | Y | Y | P | P | Y | Y | N | + |
|  | Prep | Y | Y | Y | P | Y | Y | Y | Y | + |
| CFSAN051563 | XT | P | Y | Y | P | Y | Y | Y | N | + |
|  | Prep | Y | Y | Y | P | Y | Y | Y | Y | + |
| CFSAN051566 | XT | Y | Y | Y | P | P | Y | Y | Y | + |
|  | Prep | Y | Y | Y | P | Y | Y | Y | Y | + |
| CFSAN051758 | XT | Y | Y | Y | P | Y | Y | Y | N | + |
|  | Prep | Y | Y | Y | P | P | Y | Y | Y | + |
| CFSAN051761 | XT | Y | Y | Y | P | P | Y | Y | N | + |
|  | Prep | Y | Y | P | P | P | Y | P | Y | + |
| CFSAN051762 | XT | Y | Y | Y | P | P | Y | Y | N | + |
|  | Prep | Y | Y | Y | P | P | Y | Y | Y | + |
| CFSAN051763 | XT | Y | P | Y | P | P | P | Y | N | + |
|  | Prep | Y | Y | Y | P | Y | Y | Y | Y | + |
| CFSAN051765 | XT | Y | Y | Y | P | Y | Y | Y | P | + |
|  | Prep | Y | Y | Y | P | Y | Y | Y | Y | + |
| CFSAN051766 | XT | Y | Y | Y | P | Y | Y | Y | Y | + |
|  | Prep | Y | Y | Y | P | Y | Y | Y | Y | + |
| CFSAN051769 | XT | Y | Y | Y | P | Y | Y | Y | P | + |
|  | Prep | Y | Y | Y | P | P | Y | Y | Y | + |
| CFSAN051772 | XT | Y | Y | Y | P | P | Y | Y | P | + |
|  | Prep | Y | Y | Y | P | Y | Y | Y | Y | + |
| CFSAN052204 | XT | Y | Y | Y | P | P | Y | Y | P | + |
|  | Prep | Y | Y | Y | P | Y | Y | Y | Y | + |
| CFSAN052205 | XT | Y | Y | Y | P | P | Y | Y | P | + |
|  | Prep | Y | Y | Y | P | P | Y | Y | Y | + |
| CFSAN052206 | XT | Y | Y | Y | P | P | Y | Y | P | + |
|  | Prep | Y | Y | Y | P | Y | Y | Y | Y | + |
| CFSAN052207 | XT | Y | Y | Y | P | P | Y | Y | P | + |
|  | Prep | Y | Y | Y | P | P | Y | Y | Y | + |
| CFSAN051559 | XT | Y | Y | Y | P | Y | N | N | N | - |
|  | Prep | Y | Y | Y | P | P | N | N | N | - |
| CFSAN051561 | XT | Y | Y | Y | P | P | N | N | N | - |
|  | Prep | Y | Y | Y | P | P | N | N | N | - |
| CFSAN051562 | XT | Y | Y | Y | Y | P | N | N | N | - |
|  | Prep | Y | Y | Y | Y | Y | N | N | N | - |
| CFSAN051564 | XT | Y | Y | Y | P | P | N | N | N | - |
|  | Prep | Y | Y | Y | P | P | N | N | N | - |
| CFSAN051565 | XT | Y | Y | Y | P | P | N | N | N | - |
|  | Prep | Y | Y | Y | P | Y | N | N | N | - |
| CFSAN051567 | XT | Y | Y | Y | P | P | N | N | N | - |
|  | Prep | Y | Y | Y | P | P | N | N | N | - |
| CFSAN051568 | XT | Y | Y | Y | P | Y | N | N | N | - |
|  | Prep | Y | Y | Y | P | Y | N | N | N | - |
| CFSAN051759 | XT | Y | Y | Y | P | P | N | N | N | - |
|  | Prep | Y | Y | Y | P | Y | N | N | N | - |
| CFSAN051760 | XT | Y | Y | Y | P | Y | N | N | N | - |
|  | Prep | Y | Y | Y | P | P | N | N | N | - |
| CFSAN051764 | XT | Y | Y | Y | P | P | N | N | N | - |
|  | Prep | Y | Y | Y | P | P | N | N | N | - |
| CFSAN051767 | XT | Y | Y | Y | Y | P | N | N | N | - |
|  | Prep | Y | Y | Y | P | Y | N | N | N | - |
| CFSAN051768 | XT | Y | Y | Y | P | P | N | N | N | - |
|  | Prep | Y | Y | Y | P | P | N | N | N | - |
| CFSAN051770 | XT | Y | Y | Y | P | P | N | N | N | - |
|  | Prep | Y | Y | Y | P | Y | N | N | N | - |
| CFSAN051771 | XT | Y | Y | Y | P | P | N | N | N | - |
|  | Prep | Y | Y | Y | P | Y | N | N | N | - |
| CFSAN052208 | XT | Y | Y | Y | P | Y | N | N | N | - |
|  | Prep | Y | Y | Y | P | Y | N | N | N | - |

Genes in bold are found on the plasmid. Y = Identified, N = Not identified, P = Partial identification.
All strains were positive for *eae*, *stx2a*, *espA*, *espB*, *espJ*, *espK*, *fdeC*, *lpfA*, *nleA*, *nleB*, and *tir*.
<sup>a</sup>PCR specific targeting the *exhA* gene in the plasmid. If present + and not present -.

In the case of the virulence genes, AMRFinder plus identified that these strains carried intimin subtype epsilon and *stx2a* in all assemblies (Table 3). For those virulence genes found on the chromosome, there were two genes (*nleC* and *tccP*) that could only be partially identified a majority of the time, regardless of library preparation. The main difference in virulence gene detection was in the genes found on the plasmid (*exhA, espP, toxB*).

Interestingly some strains (n = 18) irrespective of the library preparation method used, resulted in the lack of presence of all three virulence genes found in the plasmid. This oddity could be due to those strains lacking the virulence plasmid entirely. To verify our hypothesis, that these strains either may have lost the plasmid during serial culture in the laboratory or if two populations (one with the plasmid and one without) were originally present, a previously reported PCR screen for detecting the *exhA* gene was used (14). Some of the strains that showed a positive result for *exhA* by *in silico* analysis, as well as the closed strain FNW19M81 were used as the positive controls in the PCR (14). The *exhA* positive strains resulted in a PCR product with the correct size (∼1500 bp) while the *exhA* negative strains by *in silico* WGS analyses resulted in no PCR product. The lack of *exhA* PCR products on the 18 strains combined with the results from the *in silico* analysis of the other two plasmid virulence genes (*espP* and *toxB*) confirmed the lack of the plasmid in these strains.

For those strains that carried the plasmid (n=15), XT libraries identified *exhA* in all strains (93.3% complete and 6.7% partial at 55.5% gene coverage), *espP* in all strains (100% complete), and *toxB* in nine strains (13.3% complete and 46.7% partial at an average of 56.6% gene coverage). DNA Prep libraries had higher rates of identification of the plasmid genes: *exhA* in all strains (100% complete), *espP* in all strains (93.3 % complete and 6.7% partial at 56.5% gene coverage), and *toxB* in all strains (100% complete). The assemblies and raw reads from XT libraries did not cover the entire *toxB* gene unlike those from DNA Prep as can be seen in Figure 3.

**Figure 3.**
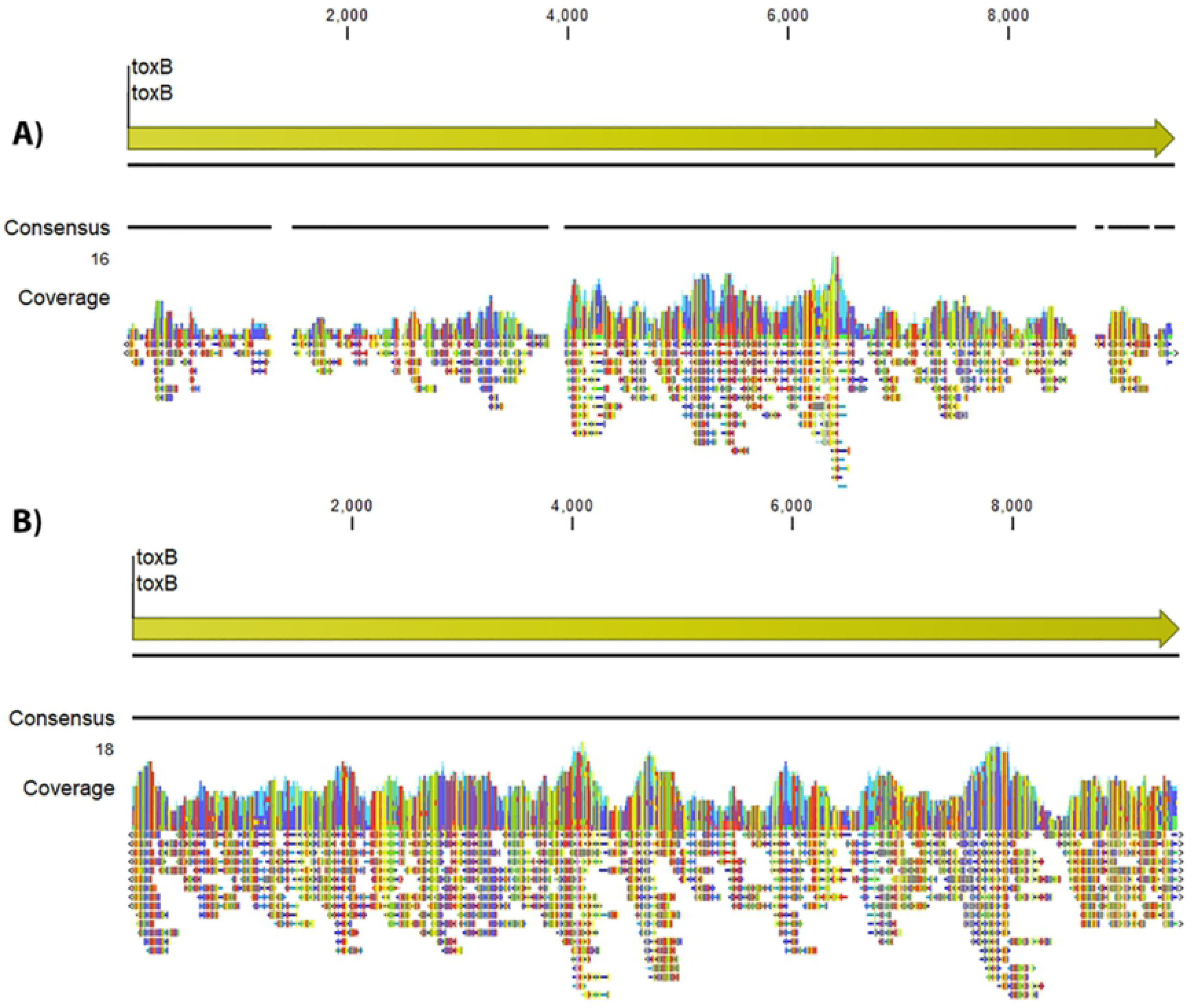
Reference mapping of the *toxB* gene (we used the *toxB* gene sequence from strain Sakai O157:H7 as reference). A) Nextera XT output for strain FNW19M81 showing that the reads did not cover the entire gene and therefore that gene was not present in the final XT assembly. B) Nextera DNA Prep output for the same strain showing that it covered the entire *toxB* gene and therefore that gene was present in the final Prep assembly.

### SNP phylogenetic analysis

To test that both library preparations (Prep vs XT) produced the same phylogenetic result (SNP calls), the CFSAN SNP pipeline (27) was used to analyze the raw data for all strains from both library preparations using FNW19M81 as the reference and three near-neighbor clinical isolates that were unrelated to the outbreak as outliers. The maximum likelihood tree generated by the SNP matrix is shown in Figure 4. All 30 of the flour isolates clustered together and were separated by 0 – 2 SNPs. The three unrelated clinical isolates were separated by 73 – 130 SNPs from the flour cluster. There were no SNP differences identified between the different library preparation methods.

**Figure 4.**
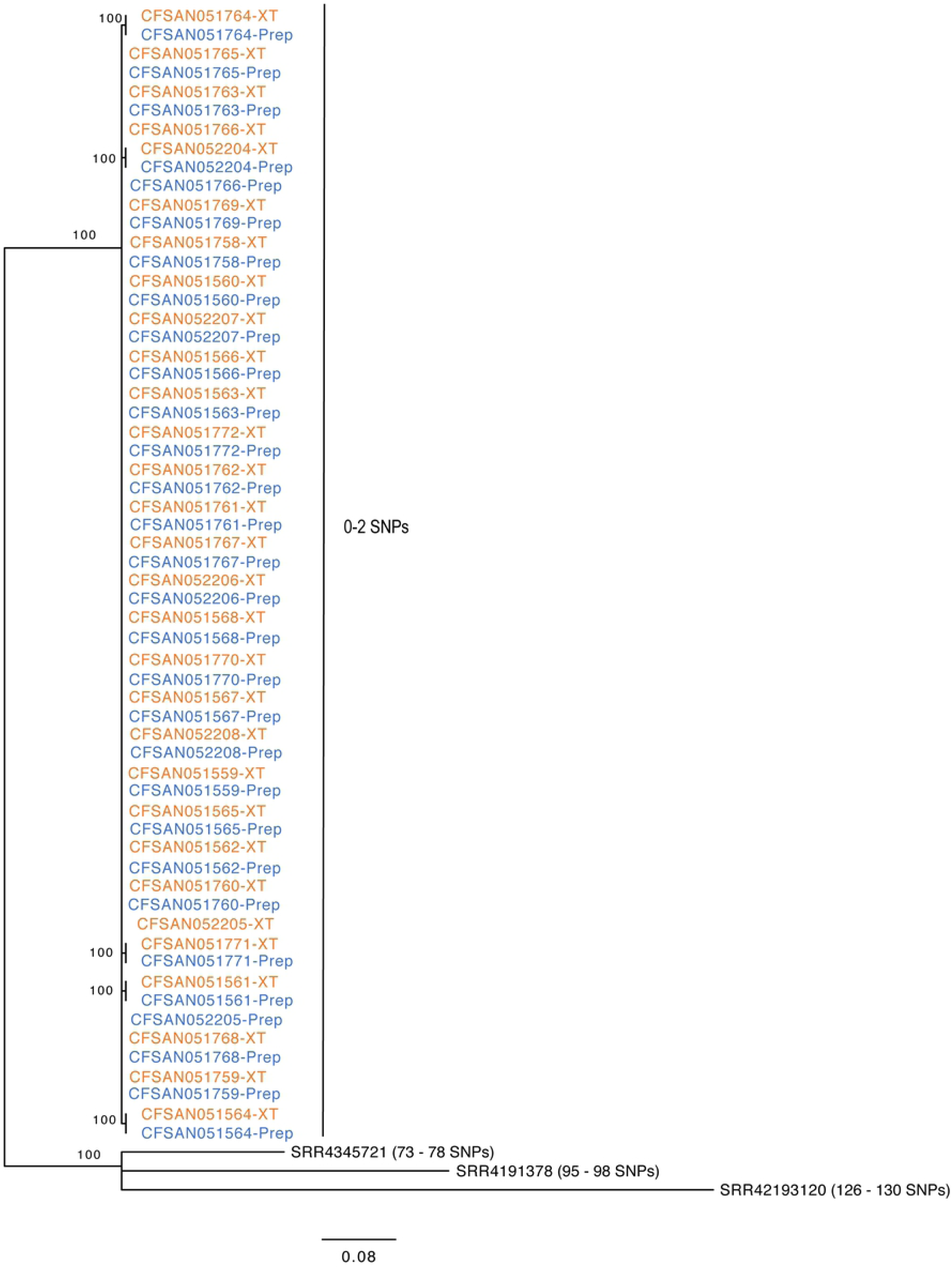
Phylogenetic tree obtained by a maximum-Likelihood analysis of the resultant SNP matrix from the CFSAN SNP pipeline of the data from the 30 strains obtained from each DNA library kit. Results obtained with the Nextera XT libraries are shown in red while DNA Prep libraries are show in blue. The genome of FNW19M81 was used as the reference.

## Discussion

Whole genome sequencing is comprised of many different modules and each of those may influence the end results (29, 30). There are two modules, bacterial culture and DNA extraction, prior to library preparation and any artifacts from those methods have the potential to affect the resultant library and sequence data. Therefore, in this study, we used the same DNA extracts to directly compare the data results obtained from the two different DNA library preparations (XT and DNA Prep). Comparison of the two DNA library preparation kits, with the same DNA extracted from the 2016 outbreak flour strains, shows that the DNA Prep kit outperforms the XT kit in terms of both quality of the data generated and genome coverage. The importance of complete WGS of *E. coli* cannot be understated for public health importance, making the library preparation method extremely crucial to ensure complete and accurate sequencing. The results of this study correlate with the findings of previous studies using the DNA Prep method, where in the bead-linked transposome allows for improved coverage uniformity in difficult to sequence regions (2, 4), and, furthermore, our study shows this kit generates a more stable coverage of the complete genome of the isolates.

One advantage of the DNA prep kit is that the transposon is linked to a bead, and the linkage of the enzyme to a bead controls the tagmentation process thus the median insert size in the resulting library. Not only were the average insert size distributions larger for DNA Prep, there is overall more stability in the quality of the libraries prepared using this kit compared to XT (335 ± 17.8 bp and 260 ± 28.8 bp, respectively). The larger, more stable insert size using DNA Prep allowed for less sequencing overlap (more bases of the genome to be sequenced), leading to better quality runs with better cluster density, depth of sequencing, and overall better assemblies. MiSeq output for DNA Prep runs showed Q30 read qualities above 90% in comparison to XT runs which had Q30 qualities at 74%. Overall, the MiSeq run quality for libraries prepared with DNA Prep had higher Q30%, higher PF%, and stable cluster density when compared with the runs of libraries prepared with XT. This impact can be seen in the average read length at or above Q30; higher Q30 average length leads to better quality assemblies and this metric reflects the difference in overall quality of Prep vs XT. These factors combined strongly indicate the robust and consistent nature of the DNA Prep library preparation. Additionally, Prep reactions were simple to setup, takes roughly 4 hours to complete with equivalent to or less hands-on time than XT, and the DNA Prep kit allows for greater customization of the target insert size. The combination of larger insert size, higher Q30 and better quality reads results in less run failures and more accurate assemblies (fewer contigs with higher N50 length) for libraries prepared with DNA Prep, which shows the value of this library preparation kit.

The identification of virulence genes during outbreaks is critical for discrete characterization of the type and pathogenesis of the *E. coli* pathogen. There have been previous outbreaks, such as the 2011 *E. coli* O104:H4 centered in Germany but affecting various other countries in the European Union (31, 32) where the strain exhibited a unique combination of virulence genes. The virulence genes present resulted in a strain that had unusual pathogenicity and outcomes in human illnesses, leading to higher proportion of HUS cases (∼23%) compared to classic HUS rates for STECs (6%) (33). This O104:H4 strain belonged to ST678, and produced Stx2a but whole-genome sequencing (WGS) revealed that it was 93% identical to enteroaggregative *E. coli* (EAEC) strain 55589 (33-35). Taken together the genetic analyses revealed that this strain was an EAEC strain that had acquired the ability to produce Stx via phage conversion. WGS is becoming the gold standard in surveillance of foodborne pathogens and it is vital to identify all virulence genes present, in order to make sound public health decisions.

The method of library preparation impacts the ability to identify all virulence genes, including those which present unique genomic challenges. The aforementioned 2011 outbreak emphasizes the necessity for a library preparation method which is able to capture the entire genome in the data. In this study, one of the largest discrepancies between library preparations was seen regarding the *toxB* gene. The *toxB* is a 10-kb virulence gene that is distributed among enterohemorrhagic *E. coli* (EHEC) and enteropathogenic *E. coli* (EPEC) (36). The assemblies from XT identified this gene 58% of the time compared to 100% identification from DNA Prep assemblies. It is possible that due to the large size of the gene, the low GC content (31%), as well as being surrounded by insertion sequences, makes identification of this gene more difficult. The ability for DNA Prep to sequence problematic regions has been previously shown in organisms with wide ranges of GC content (4) and that the fragmentation step using the bead-linked transposome allows for more even coverage across the entire genome. Additionally, plasmids can be difficult to capture with WGS, but we believe the increased amount of DNA input coupled with the lack of bias in the fragmentation allow for more of the plasmid to be sequenced. Identification of plasmid-borne genes is a necessity due to the increase in antimicrobial resistant plasmids and the ability for bacteria to acquire them via uptake or horizontal transfer. The DNA Prep library kit allows these plasmids to be sequenced, increasing the ability for public health surveillance of novel and critical AMR genes.

The results from this preliminary data set illustrates the benefit of using DNA Prep library preparation as replacement for the XT preparation for virulence typing in *E. coli*. Coverage of the plasmid is enhanced due to higher DNA input and overall coverage of the chromosome is evenly distributed. Virulence genes were identified at a higher rate and overall run quality was improved by utilizing the DNA Prep library preparation method. Overall, the DNA Prep results showed more consistent insert size regardless of bacterial species, GC content, or DNA input which was not observed with XT.

